# Complex plant metabolomes guide fitness-relevant foraging decisions of a specialist herbivore

**DOI:** 10.1101/2020.07.13.200618

**Authors:** Ricardo A. R. Machado, Vanitha Theepan, Christelle A.M. Robert, Tobias Züst, Lingfei Hu, Qi Su, Bernardus C. J. Schimmel, Matthias Erb

## Abstract

Plants produce complex mixtures of primary and secondary metabolites. Herbivores use these metabolites as behavioral cues to increase their fitness. However, how herbivores integrate different metabolite classes into fitness-relevant foraging decisions *in planta* is poorly understood. We developed a molecular manipulative approach to modulate the availability of sugars and benzoxazinoid secondary metabolites as foraging cues for a specialist maize herbivore, the western corn rootworm. By disrupting sugar perception in the western corn rootworm and benzoxazinoid production in maize, we show that sugars and benzoxazinoids act as distinct and dynamically integrated mediators of short-distance host finding and acceptance. While sugars improve the capacity of rootworm larvae to find a host plant and to distinguish post-embryonic from less nutritious embryonic roots, benzoxazinoids are specifically required for the latter. Host acceptance in the form of root damage is increased by benzoxazinoids and sugars in an additive manner. This pattern is driven by increasing damage to post-embryonic roots in the presence of benzoxazinoids and sugars. Benzoxazinoid- and sugar-mediated foraging directly improves western corn rootworm growth and survival. Interestingly, western corn rootworm larvae retain a substantial fraction of their capacity to feed and survive on maize plants even when both classes of chemical cues are almost completely absent. This study unravels fine-grained differentiation and integration of primary and secondary metabolites into herbivore foraging and documents how the capacity to compensate for the lack of important chemical cues enables a specialist herbivore to survive within unpredictable metabolic landscapes.

## Introduction

Herbivore foraging behavior contributes to the distribution and performance of herbivores and plants in natural and agricultural ecosystems (1–3). Insect herbivores often exhibit pronounced oviposition and feeding preferences for specific plant species, genotypes within species physiological states within genotypes (1, 4, 5). Most insect herbivores also show characteristic preferences for specific plant organs and tissues (6–8).

Herbivores establish preferences through different types of host-cues (9–12). Chemical cues, including plant primary and secondary metabolites, are particularly important for herbivores, as they provide specific information about the identity, physiological status and nutritional value of host plants and tissues (13–15). The overarching view in the field of chemical ecology is that primary metabolites are used as cues to identify nutritious hosts and tissues, while volatile and non-volatile secondary metabolites are used as indicators of toxicity and defense status, and as signature cues of specific host plant lineages and species (16–18). Specialized herbivores in particular are often attracted and stimulated by host-specific secondary metabolites (19). Over the past decades, herbivores have been found to respond to a multitude of plant primary and secondary metabolites in artificial diet experiments (13, 16, 17). An increasing number of studies now also document the importance of these metabolites *in planta* through molecular manipulative approaches (20–26).

Plants produce diverse sets of primary and secondary metabolites (27), and herbivores likely integrate many of these metabolites into their foraging behavior (16, 28, 29). Chemical cue integration is thought to allow herbivores to obtain more accurate information about the nutritional value and toxicity of complex host plant metabolomes (17) and thus to increase the robustness of their foraging decisions (12, 29). Many insect herbivores are known to be attracted to specific combinations of volatile chemicals, for instance (14, 29). Furthermore, some herbivores avoid combinations of secondary metabolites more strongly than individual compounds (30–32). A limited number of studies also indicate that herbivores may be able to integrate primary and secondary metabolites into their foraging strategies (17). Using artificial diet experiments, it was found that tannins reduce food intake by locusts at low protein:carbohydrate ratios, but not at high ratios, a behavior which mirrored the conditional impact of tannins on locust performance (33, 34). Modulation of prior exposure to secondary metabolites on subsequent food choice was observed for both insect herbivores (35) and mammals (36, 37). Conversely, sugars were found to mask the aversive taste of secondary metabolites (38). Despite these advances, we currently lack a detailed understanding of how primary and secondary metabolites interact to determine herbivore behavior under biologically realistic conditions (9). The paucity of manipulative experiments that test for interactions between host-derived chemical foraging cues *in planta* limits our capacity to assess the concerted impact of different metabolites on herbivore feeding preferences, and, more generally, our understanding of the role of plant metabolic complexity in herbivore behavior and plant-herbivore interactions.

An implicit assumption of herbivore foraging theory is that herbivore behavior improves herbivore fitness (39). With some notable exceptions, herbivores generally prefer to oviposit and feed on plants and tissues that increase their performance (40–42). Host plant chemicals likely play an important role in these preference-performance relationships (12, 17). However, the contribution of individual behavioral cues to herbivore fitness in the context of the full chemical complexity of a given host plant has remained difficult to quantify (30, 43, 44). Targeted molecular manipulation of the production and perception of chemical cues provides new opportunities in this context, including the evaluation of the benefits of the integration of multiple chemical cues into herbivore foraging (29), and the assessment of the importance of the flexible use of multiple foraging cues with redundant information content (11, 12).

Insect herbivores use chemosensory receptors to detect volatile and non-volatile plant chemicals (45, 46). Gustatory receptors (GRs) are required for the detection of a variety of chemicals (47–49), including non-volatile plant chemicals such as alkaloids (24) and sugars (50). Knocking out specific GRs reduces oviposition of swallowtail (*Papilio xuthus*) butterflies (24) and host recognition by silkworm (*Bombyx morĩ*) larvae (23), thus demonstrating the functional importance of individual GRs for herbivore behavior. Work in Drosophila (*Drosophila melanogaster*) revealed that some GRs are broadly tuned and can mediate avoidance to many different compounds (51), while others are narrowly tuned and confer responsiveness to specific compounds (52). Gr43a is a highly conserved insect taste receptor that specifically responds to D-fructose in Drosophila (49), the diamondback moth (*Plutella xylostella*) (53) and the cotton bollworm (*Helicoverpa armigera*) (54). As Gr43a is responsible for the detection D-fructose in the hemolymph as a proxy for carbohydrate supply, *Gr43a* silenced flies also become unresponsive to other dietary sugars (55). Interestingly, different GRs can interact dynamically through competition, inhibition and activation (47, 56, 57), resulting in substantial potential for GR-mediated integration of multiple chemical cues into behavioral responses.

Here, we developed a manipulative approach to evaluate the importance of maize primary and secondary metabolites for the foraging and foraging-dependent performance of the western corn rootworm (*Diabrotica virgifera virgifera*). The western corn rootworm is an economically damaging maize pest (58). Its larvae are highly specialized on maize roots (59). Over the last decades, the chemical ecology of the western corn rootworm has been studied in detail (13, 58, 59). Field and laboratory studies revealed tight associations between maize root chemistry and western corn rootworm behavior. Western corn rootworm larvae respond behaviorally to a wide variety of chemical cues, including CO_2_ (60), sugars and fatty acids (61, 62), aromatic and terpene volatile organic compounds (63, 64), conjugated phenolic acids (65) and benzoxazinoids (66). The emerging picture is that these chemicals likely allow western corn rootworm larvae to locate plant roots from a distance (67), to discriminate between plants of different quality (63–65, 68), and to identify and feed on the most nutritious roots (66, 69). Several *in vitro* experiments also suggest that the western corn rootworm can integrate multiple chemical cues for host finding and acceptance (61), hinting at the substantial sensory capacity of this specialist root feeder. By independently manipulating the availability of benzoxazinoids and sugars as foraging cues of the western corn rootworm through plant genetics and insect RNA interference (RNAi), we demonstrate that sugars and benzoxazinoids serve both specific and integrated roles as determinants of the behavior and behaviorally driven performance of this specialist herbivore.

## Results

### The western corn rootworm prefers to feed on root tissues that are rich in benzoxazinoids and soluble sugars

The root system of young maize plants consists of embryonic and post-embryonic roots (Fig. 1*A*). Embryonic roots emerge directly from the embryo and comprise primary and seminal roots. Post-embryonic roots emerge from the hypocotyl and stem and comprise crown roots, and, at later developmental stages of the plant, internode-derived brace roots (70). To determine feeding preferences of the western corn rootworm within the root system of young maize plants, we infested soil-grown maize plants with western corn rootworm larvae for 7 days and then scored the damage on the different root types. In line with earlier studies (66, 69), we found low amounts of damage on embryonic roots and substantial damage on post-embryonic roots (Fig. 1*B*). While embryonic roots showed scattered bite marks, post-embryonic roots were often partially or even fully removed (Fig. S1). To test whether this feeding preference is reflected in the distribution of the larvae within the root system, we carried out a series of behavioral experiments. First, we laid out intact maize root systems on a filter paper and then recorded the position of second instar larvae at different time points after their release. Five times more larvae were found on post-embryonic than embryonic roots 30 min after the release of the larvae (Fig. 1*C*). This preference persisted over the duration of the experiment. To test whether the preference of the larvae for post-embryonic roots may be due to differences in abundance or the relative position of the two root types within the root system (Fig. 1*A*), we offered post-embryonic and embryonic root pieces of equal size to the larvae. Similar to what was observed for entire root systems, significantly more larvae were found on the post-embryonic root pieces than the embryonic root pieces after 30 min (Fig. 1*D*). Thus, the western corn rootworm preferentially stays and feeds on post-embryonic roots of young maize plants.

**Fig. 1.**
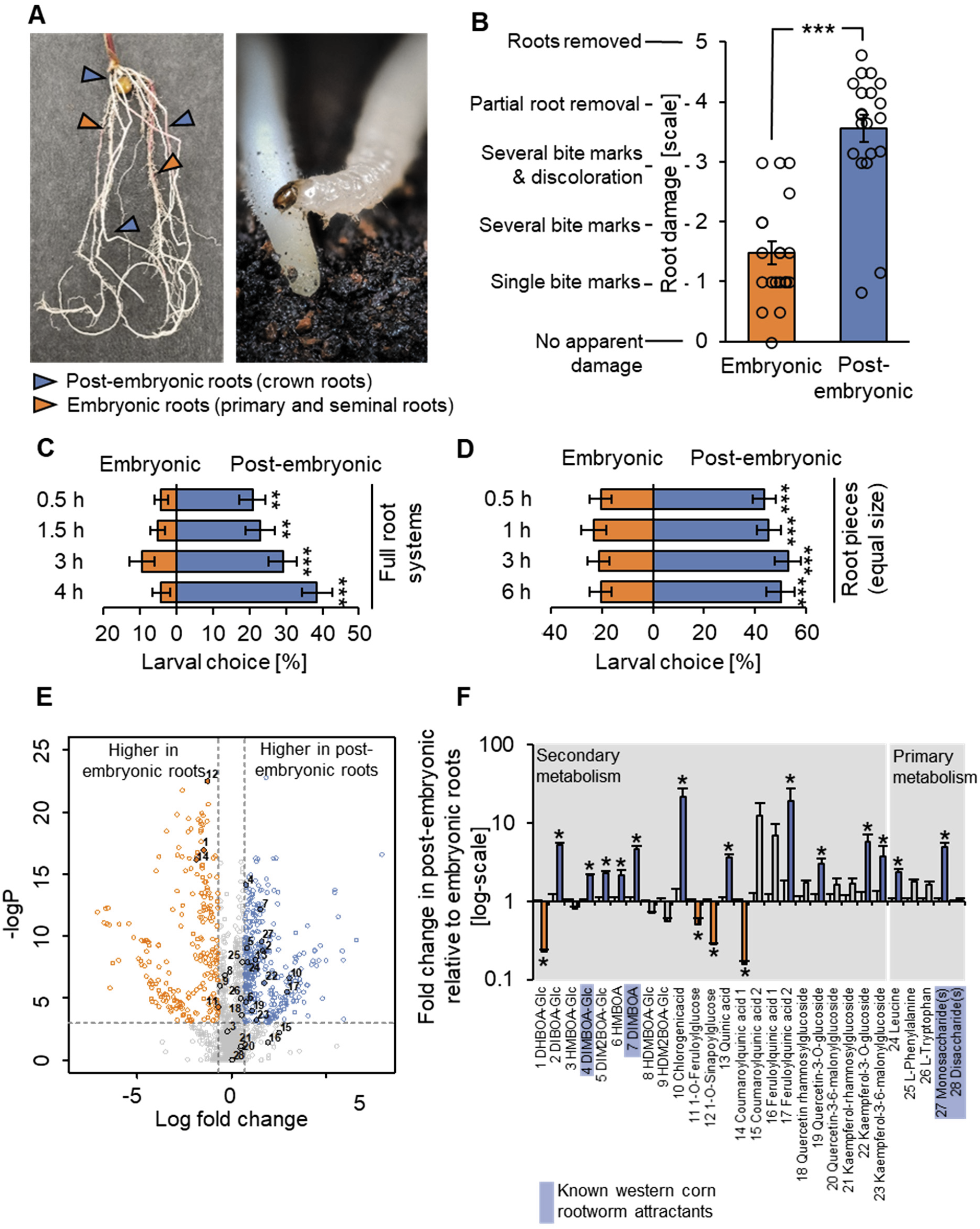
The western corn rootworm prefers to feed on root tissues that are rich in benzoxazinoids and soluble sugars. (*A*) Left: young maize plants produce embryonic primary and seminal roots (orange arrows) and post-embryonic crown roots (blue arrows). Right: Western corn rootworm larvae are highly specialized maize root feeders that are very mobile in the second and third instar. A third instar larvae is shown. (*B*) Average damage score observed on post-embryonic and embryonic roots of soil-grown maize plants after 7 days of infestation by western corn rootworm larvae (***p<0.001, Wilcoxon Signed Rank test, n=20 plants with 15 larvae each, data from Fig. 4*D*). Dots represent damage scores on individual plants (averaged within root types). For frequency distributions of individual roots, refer to Fig. S1. (*C*) Preference of western corn rootworm larvae for post-embryonic and embryonic roots within the root system (**p<0.01, ***p<0.001, FDR-corrected Least Square Mean post hoc tests, n=16 dishes with 6 larvae each). (*D*) Preference of western corn rootworm larvae for root pieces of equal size of post-embryonic and embryonic roots (***p<0.001; FDR-corrected Least Square Mean post hoc tests, n=18 dishes with 5 larvae each). (*E*) Metabolomics profiles of methanolic extracts of post-embryonic and embryonic roots. Orange features are more abundant in embryonic roots, blue features are more abundant in post-embryonic roots (min. 2-fold difference, p<0.05, FDR-corrected Student’s *T*-tests, n=9-10). Numbers denote features that were tentatively assigned to structures based on exact mass and fragment information (Table S2). (*F*) Relative abundance differences of tentatively identified metabolites between post-embryonic and embryonic roots (fold change of peak areas; *p<0.05, FDR-corrected Student’s *T*-tests, n=9-10). Picture credits: Ricardo Machado, Lingfei Hu, Cyril Hertz. Error bars denote standard errors of means (SEM).

To gain insights into the phytochemical differences that may drive the preference of the western corn rootworm for post-embryonic roots, we performed untargeted metabolomics on methanolic extracts of both root types. Using UHPLC-Q-TOF-MS and ESI-, 4512 mass features were detected, 358 and 250 of which were enriched in post-embryonic or embryonic roots based on more than two-fold difference in normalized signal intensity at a statistical significance of p<0.05 (Fig. 1*E*). Using exact masses and fragmentation patterns, 28 metabolites could be putatively identified within the dataset. Identified metabolites included benzoxazinoids, phenolic acid derivatives, amino acids and sugars (Fig. 1*F*, table S1). Seventeen of these metabolites showed root-type specific accumulation patterns (Fig. 1*F*). Among the secondary metabolites, benzoxazinoids and phenolic acids showed pronounced shifts in their profiles, with some metabolites accumulating in higher amounts in post-embryonic roots and others accumulating in higher amounts in embryonic roots (Fig. 1*F*). Among the detected primary metabolite features, amino acids and sugars were more abundant in post-embryonic-than in embryonic roots, with the most pronounced shifts observed for features matching leucin and monosaccharides (Fig. 1*F*). Thus, the preference of the western corn rootworm for post-embryonic roots is associated with distinct primary and secondary metabolite accumulation patterns in these tissues.

To narrow down the list of metabolites that may prompt the western corn rootworm to feed on post-embryonic roots, we performed a literature search on western corn rootworm attractants and feeding stimulants. Thirteen different compounds known to elicit behavioral responses in the western corn rootworm *in vitro* were found (Table S2). Cross-referencing this table with the list of chemicals that accumulated in post-embryonic roots at higher concentrations resulted in the identification of four candidate compounds that may mediate post-embryonic root preference: The monosaccharides glucose and fructose, which act as feeding stimulants (61) and the benzoxazinoids DIMBOA-Glc and DIMBOA, which form iron complexes in the rhizosphere that act as short-distance host preference and acceptance cues (66). Based on these results and earlier studies (66, 69), we hypothesized that benzoxazinoids and monosaccharides may interact to determine the preference of the western corn rootworm for post-embryonic roots. Given the observed overlap between post-embryonic root preference and overall plant attractiveness (66), we also hypothesized that the same compounds may mediate general host recognition and acceptance in this specialist herbivore.

### Independent manipulation of benzoxazinoids and sugars as foraging cues

To test for the individual and combined roles of benzoxazinoids and sugars in mediating western corn rootworm behavior, we sought to independently manipulate their availability as foraging cues. The first dedicated step in benzoxazinoid biosynthesis is mediated by the indole-3-glycerol phosphate lyase *Bx1* (71). By consequence, *bx1* mutants are benzoxazinoid deficient (71, 72). To test if *bx1* mutant plants can be used to manipulate benzoxazinoids independently of root sugars, we performed a series of targeted analyses of wild type (WT) B73 and *bx1* roots. In accordance with our metabolomics screen and earlier studies (66, 69), post-embryonic roots of WT plants contained 4-fold higher DIMBOA levels and 2-to 3-fold higher DIMBOA-Glc and DIM_2_BOA-Glc levels, but lower HDMBOA-Glc levels than embryonic roots (Fig. 2*A*). Benzoxazinoid levels in the *bx1* mutant were reduced by more than 97%, and residual benzoxazinoid concentrations were not significantly different between crown and primary roots (Fig. 2*A*). As expected from the metabolomic analyses, post-embryonic roots of WT plants contained higher levels of the monosaccharides glucose and fructose than embryonic roots (Fig 2*A*). Sucrose levels were also higher in post-embryonic roots (Fig 2*A*). Sugar profiles were similar in WT and *bx1* roots (Fig. 2*A*), demonstrating that benzoxazinoid availability can be manipulated independently of sugar availability. The differential abundance between and crown and primary roots of the other 17 identified metabolic features was conserved in the *bx1* mutant as well (Fig. S2). We only detected a significant interaction between root type and genotype for tryptophan, which is also derived from indole-3-glycerol phosphate (71). Tryptophan accumulated in higher amounts in post-embryonic roots of both wild type and *bx1* mutant plants, and this difference was more pronounced in the *bx1* mutant (Fig. S2). Thus, we conclude that apart from benzoxazinoids, the metabolic differences between crown and primary roots are conserved in the *bx1* mutant, thus validating the use of this genotype to assess the behavioral impact of benzoxazinoids on western corn rootworm behavior.

**Fig. 2.**
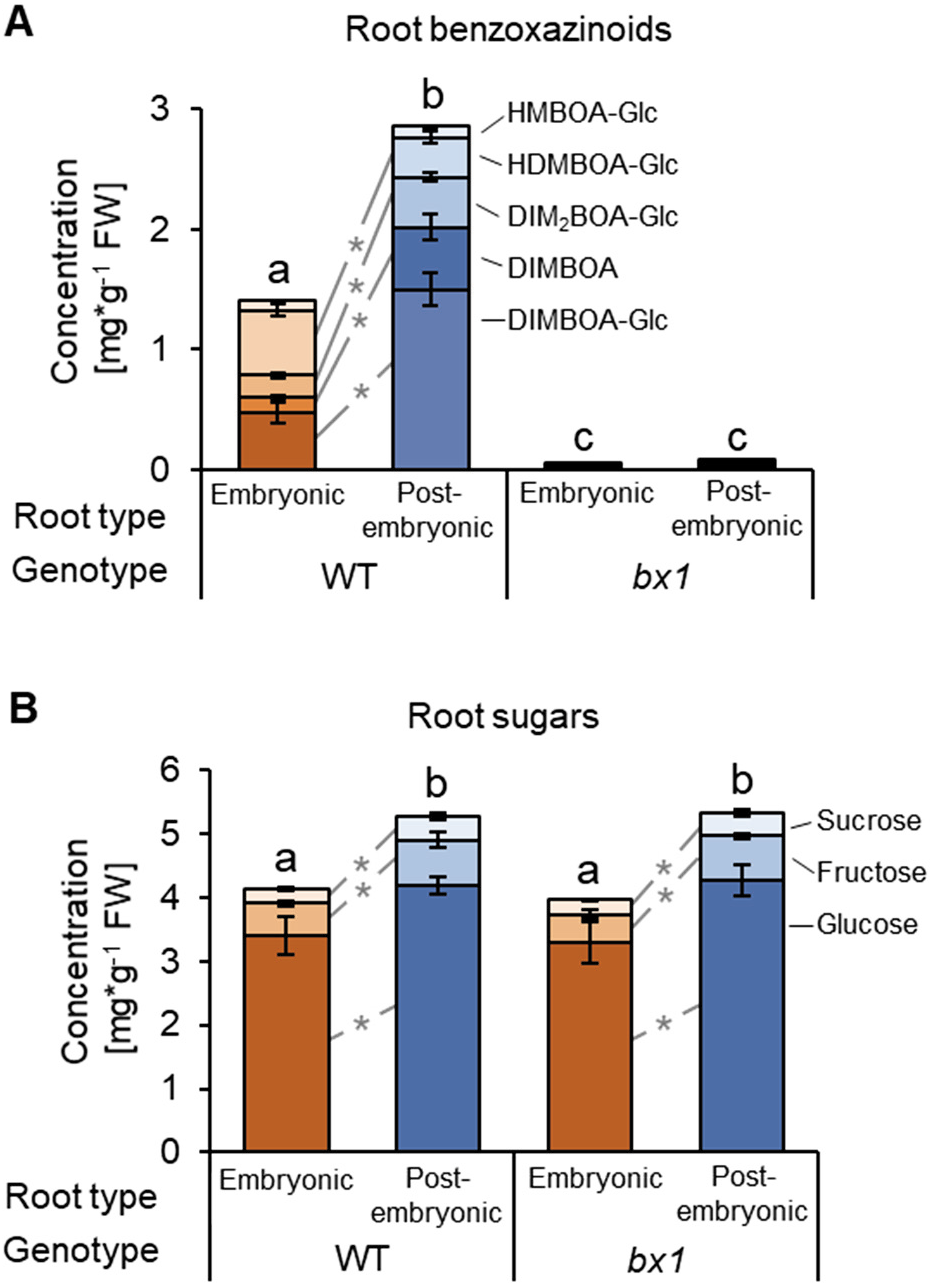
A mutation in the Bx1-gene suppresses root-type specific benzoxazinoid accumulation independently of root sugars. (*A*) Concentrations of benzoxazinoids in embryonic and post-embryonic roots of wild type B73 and *bx1* mutant plants (n=11-18). (B) Concentrations of glucose, fructose and sucrose in embryonic and post-embryonic roots of wild type B73 and *bx1* mutant plants (n=8). Letters indicate significant differences in total amounts between root types and genotypes (p<0.05, Holm-Sidak post hoc tests). Asterisks indicate significant differences in the concentrations of individual compounds between post-embryonic and embryonic roots (p<0.05, Holm-Sidak post hoc tests). Error bars denote standard errors of means (SEM).

To manipulate the availability of sugars as foraging cues independently of benzoxazinoids, we targeted sugar perception in the western corn rootworm. This approach was chosen because manipulation of primary metabolism in plants results in pleiotropic effects (73, 74). Sugar perception in insects is highly conserved and mediated by *Gr43a*-like genes (49, 50, 53, 54, 75–78). BLAST search of publicly available nucleotide sequences revealed a single putative *Gr43a* gene in the western corn rootworm, here named *DvvGr43a* (Fig. 3*A*). The full-length sequence of this gene can be retrieved from NCBI [Accession: XM_028279548.1]. Similarity between *DvvGr43a* and putative *Gr43a* genes from different insects, many of which have been functionally characterized, was found to be between 42% and 64% at the protein level. No other closely related genes were found in western corn rootworm genome. Structural prediction indicated that the *DvvGr43a* gene encodes a 7-transmemrane domain protein, consistent with its putative role as a gustatory receptor (Fig. 3*B*). Gene-expression profiling revealed stronger expression of *DvvGr43a* in the head than the body of western corn rootworm larvae (Fig. 3C). To manipulate *DvvGr43a*, we targeted its expression through RNA interference (RNAi) by feeding the larvae with a 240 bp dsRNA targeting *DvvGr43a*. Compared to control larvae that were fed with dsRNA targeting green fluorescent protein (GFP), this treatment resulted in a >70% reduction in *DvvGr43a* transcript abundance (Fig. 3*D*). To assess silencing specificity, we measured the transcription of the putative CO_2_-receptor *DvvGr2* (67). Feeding dsRNA targeting *DvvGr43a* did not change the transcript abundance of *DvvGr2* (Fig. 3*E*). Accordingly, the responsiveness of *DvvGr43a*-silenced larvae to volatile CO_2_ remained intact (Fig. S3). To evaluate whether *DvvGr43a* silencing changes the ability of western corn rootworm to respond to benzoxazinoids, we conducted choice assays with the behaviorally active benzoxazinoid iron complex Fe(III)(DIMBOA)3 (66). After 1 hour, the majority of western corn rootworm larvae were found on filter paper discs spiked with Fe(III)(DIMBOA)3, independently of *DvvGr43a* silencing. Next, we tested whether *DvvGr43a* silencing changes the ability of the larvae to respond to sugars. Three hours after the start of the experiment, control larvae preferred to locate on filter discs spiked with glucose, fructose, sucrose or a mixture of the three compounds (Fig. 3*G-J*). These preferences were absent in larvae fed with *DvvGr43a* dsRNA (Fig. 3 *G-J*), demonstrating that silencing *DvvGr43a* eliminates the capacity of western corn rootworm larvae to respond behaviorally to three major plant sugars. *DvvGr43a*-dependent sugar preference was similar in non-starved and starved western corn rootworm larvae (Fig. S4). Thus, combining *bx1* plant mutants and *DvvGr43a*-silenced larvae can be used to evaluate the individual and combined impact of benzoxazinoids and sugars on herbivore foraging.

**Fig. 3.**
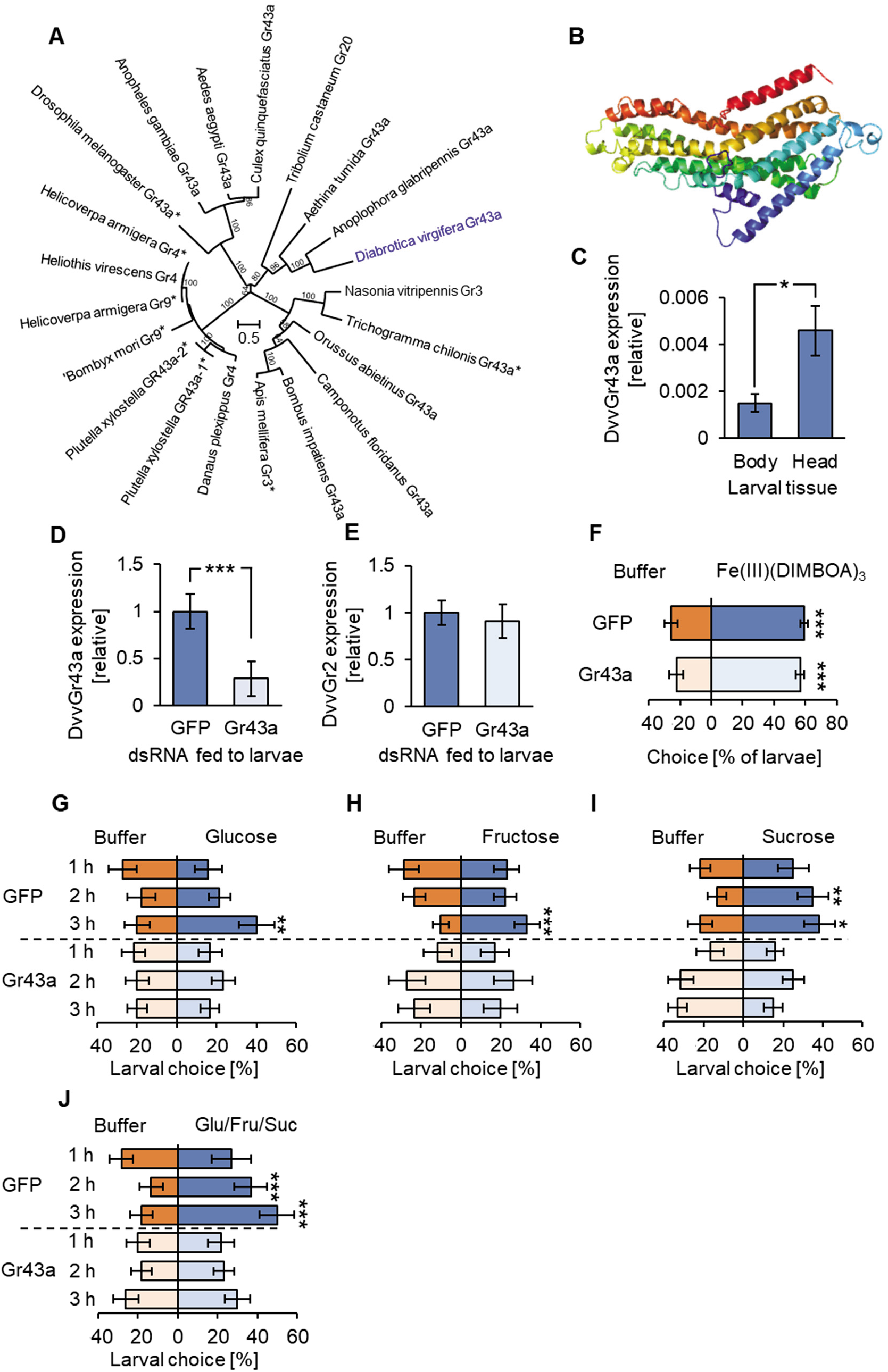
DvvGr43a mediates sugar preference of the western corn rootworm without influencing responsiveness to benzoxazinoids. (*A*) Phylogenetic relationships between gustatory sugar receptors of different insects and *DvvGr43a* of the western corn rootworm. The tree is based on protein sequences and drawn to scale, with branch lengths measured in the number of substitutions per site. Asterisks indicate functionally characterized receptors. (*B*) Protein tertiary structure of *DvvGr43a* as predicted with the Phyre2 algorithm. (*C*) *DvvGr43a* expression in the bodies (thorax and abdomen) or heads of second instar western corn rootworm larvae (*p<0.05, Student’s *T*-test n=10). (*D*) *DvvGr43a* expression in western corn rootworm larvae fed with dsRNA targeting green fluorescent protein (control, GFP) or *DvvGr43a* (Gr43a, ***p<0.001, Student’s *T*-test, n=11). (*E*) *DvvGr2* expression in western corn rootworm larvae fed with dsRNA targeting green fluorescent protein (control, GFP) or *DvvGr43a* (n=8). (*F*) Preference of GFP and *DvvGr43a* dsRNA fed larvae for buffer or Fe(III)(DIMBOA)3 on filter discs after 1 h (***p<0.001, FDR-corrected Least Square Mean post hoc tests, n=10 dishes with 6 larvae each). (*G-J*) Preference of GFP or *DvvGr43a* dsRNA fed larvae for buffer or glucose, fructose, sucrose or a mixture of the three on filter discs at different time points (*p<0.05; **p<0.01; ***p<0.001, FDR-corrected Least Square Mean post hoc tests, n=15 dishes with 6 larvae each). Error bars denote standard errors of means (SEM).

### Benzoxazinoids and sugars play distinct roles in root herbivore foraging

To understand the importance of benzoxazinoids and sugars in mediating host and tissue finding of the western corn rootworm in a neutral environment, we observed the behavior and feeding of control and *DvvGr43a*-silenced larvae on wild type and *bx1* mutant root systems laid out on moist filter paper. Ninety % of control larvae were found on the roots rather than the filter paper 4 hours after their release (Fig. 4*A*). This number dropped to 70% for *DvvGr43a* silenced larvae, independently of plant genotype. A different pattern was found when looking at the distribution of the larvae on the different root types within root systems (Fig. 4*B*). On wild type plants, significantly more control larvae were found on post-embryonic than embryonic roots. This difference was absent for control larvae feeding on *bx1* mutant plants. For *DvvGr43a* silenced larvae, more larvae were found on post-embryonic than embryonic roots, even though this preference was attenuated relative to control larvae. No preference was found for *DvvGr43a* silenced larvae feeding on *bx1* mutant roots (Fig. 4*B*). These results show that sugars contribute to root system finding and post-embryonic root preference but are not strictly required for either process. Benzoxazinoids on the other hand do not contribute to root system finding but are specifically required for the capacity of the western corn rootworm to distinguish post-embryonic from embryonic roots.

**Fig. 4.**
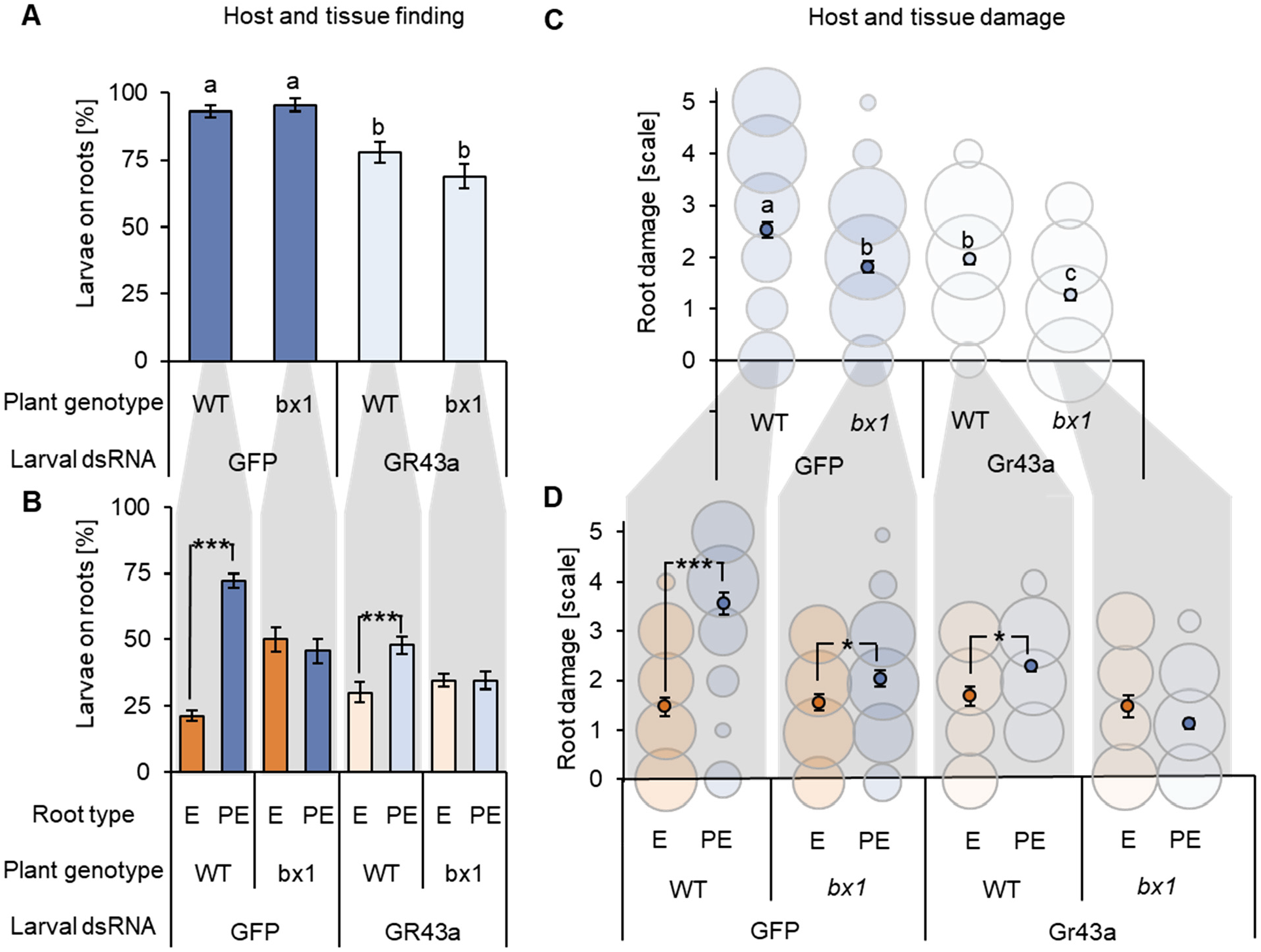
Benzoxazinoids and sugars play distinct roles in root herbivore foraging in vivo. (*A*) Proportion of control (GFP) or *DvvGr43a*-silenced (Gr43a) larvae found on the roots of wild type B73 or *bx1* mutant roots 4 hours after their release. Letters indicate statistically significant differences between treatments (Holm-Sidak post-hoc tests, p<0.05, n=15 dishes with 6 larvae each). (*B*) Proportion of control (GFP) or *DvvGr43a*-silenced (Gr43a) larvae found on embryonic (E) and post-embryonic (PE) roots within the same experiment. Asterisks indicate significant differences between root types (***p<0.001, FDR-corrected Least Square Mean post hoc tests, n=15 dishes with 6 larvae each). (*C*) Average feeding damage by control (GFP) or *DvvGr43a*-silenced larvae per root of soil-grown wild type B73 or *bx1* mutant plants 7 days after infestation. Letters indicate statistically significant differences (p<0.05, Tukey’s post hoc tests, n=20 plants with 15 larvae each). Bubble plots are shown for illustrative purposes. The sizes of the circles are proportional to the relative frequency (% within each treatment) of the different types of observed damage (each column sums up to 100%). For damage scale, refer to Fig. 1. (*D*) Average feeding damage on embryonic (E) or post-embryonic (PE) roots within the same experiment. Asterisks indicate significant differences in root damage between root types (*p<0.05, ***p<0.001 Wilcoxon Signed Rank tests, n=20 plants with 15 larvae each). Bubble plots are shown for illustrative purposes. The sizes of the circles are proportional to the relative frequency (% within each root type) of the different types of observed damage (each column sums up to 100%). Error bars denote standard errors of means (SEM).

To evaluate whether benzoxazinoids and sugars affect host acceptance in the form of active feeding, we infested roots of soil-grown wild type and *bx1* mutant plants with control or *DvvGr43a-* silenced western corn rootworm larvae and inspected the roots for characteristic feeding marks and damage after 7 days. As feeding by the western corn rootworm cannot be observed directly in the soil, and root biomass is a poor proxy for root consumption due to confounding root regrowth effects and the disproportionate impact of severed roots (79), this approach was considered most suitable for the question at hand. The experiment revealed that root damage across different roots was increased by the availability of benzoxazinoids and sugars as foraging cues in an additive fashion (Fig. 4*C*). Frequent root removal was observed in wild type plants infested with control larvae. Intermediate root damage in the form of frequent biting marks and brown discoloration was observed for *bx1* mutant roots infested with control larvae and wild type roots infested with *DvvGr43a-silenced* larvae. Infrequent biting marks and/or no visible damage was observed for *bx1* mutant roots infested with *DvvGr43a*-silenced larvae. The same additive pattern was also visible when looking at the distribution of damage patterns between the different root types (Fig. 4*D*). Post-embryonic root damage was strongest and significantly more pronounced than embryonic root damage in wild type plants infested with control larvae. Post-embryonic root damage and damage differences with embryonic roots were intermediate for *bx1* mutants infested with control larvae and wild type plants infested with *DvvGr43a*-silenced larvae. Post-embryonic root damage was lower and no longer different from damage to embryonic roots for *bx1* mutants infested with *DvvGr43a*-silenced larvae (Fig. 4*D*). Thus, both benzoxazinoids and sugars increase root damage in an additive manner, and this pattern is driven by post-embryonic root damage. Nevertheless, bite marks were still observed on 69% of *bx1* mutant roots infested with *DvvGr43a*-silenced larvae, showing that the larvae attack maize roots even in the absence of these cues. Taken together, these results reveal distinct and additive roles of benzoxazinoids and sugars in root and tissue location and feeding preferences of the western corn rootworm.

### Using benzoxazinoids and sugars as foraging cues improves herbivore growth and survival

Herbivore foraging theory predicts that herbivores use plant chemical cues to make fitness-relevant foraging decisions. To test whether the integration of primary and secondary metabolites into foraging improves herbivore performance and survival, we first evaluated whether the absence of benzoxazinoids or the inability to detect sugars may have direct performance effects on the western corn rootworm (as opposed to the effects mediated by changes in foraging behavior). Benzoxazinoids improve western corn rootworm performance under iron-limiting conditions (66) and upon attack by entomopathogenic nematodes (80, 81). When bioavailable iron is abundant and natural enemies are absent, larval performance under no-choice conditions is not altered by benzoxazinoids (66, 69). Thus, growing maize plants with bioavailable EDTA-Fe as iron source allowed us to exclude direct effects of benzoxazinoids on larval performance. To test whether silencing *DvvGr43a* has direct negative effects on western corn rootworm performance, we reared control and *DvvGr43a*-silenced larvae on young maize seedlings that produce embryonic, but no post-embryonic roots. Larval weight gain and mortality of control and *DvvGr43a-silenced* larvae after 6 days was similar (Fig. S5), demonstrating that silencing *DvvGr43a* has no direct effects on larval performance in a no-choice situation. Note that larval mortality in this experiment was >40%, which is within the normal range for early instars of this species (82, 83).

Having established the validity of our experimental approach, we proceeded to evaluate the impact of benzoxazinoid- and sugar-dependent foraging behavior on western corn rootworm performance. Highest larval weight and lowest mortality was observed for control larvae feeding on wild type plants (Fig. 5*A-B*). Intermediate weight gain and mortality was observed for control larvae feeding on *bx1* mutant roots and for *DvvGr43a*-silenced larvae feeding on wild type roots. Lowest weight gain and highest mortality was observed for *DvvGr43a-silenced* larvae feeding on *bx1* mutant roots. Larval performance thus mirrored root damage patterns (Fig. 4*C-D*). This was confirmed by correlation analysis which revealed a positive relationship between total larval weight and survival on a given plant and the total damage observed on post-embryonic roots (Fig. 5*C-F*). In conclusion, sugars and benzoxazinoids act together to improve western corn rootworm performance by acting as foraging cues that guide the herbivore to feed on the most profitable host tissue.

**Fig. 5.**
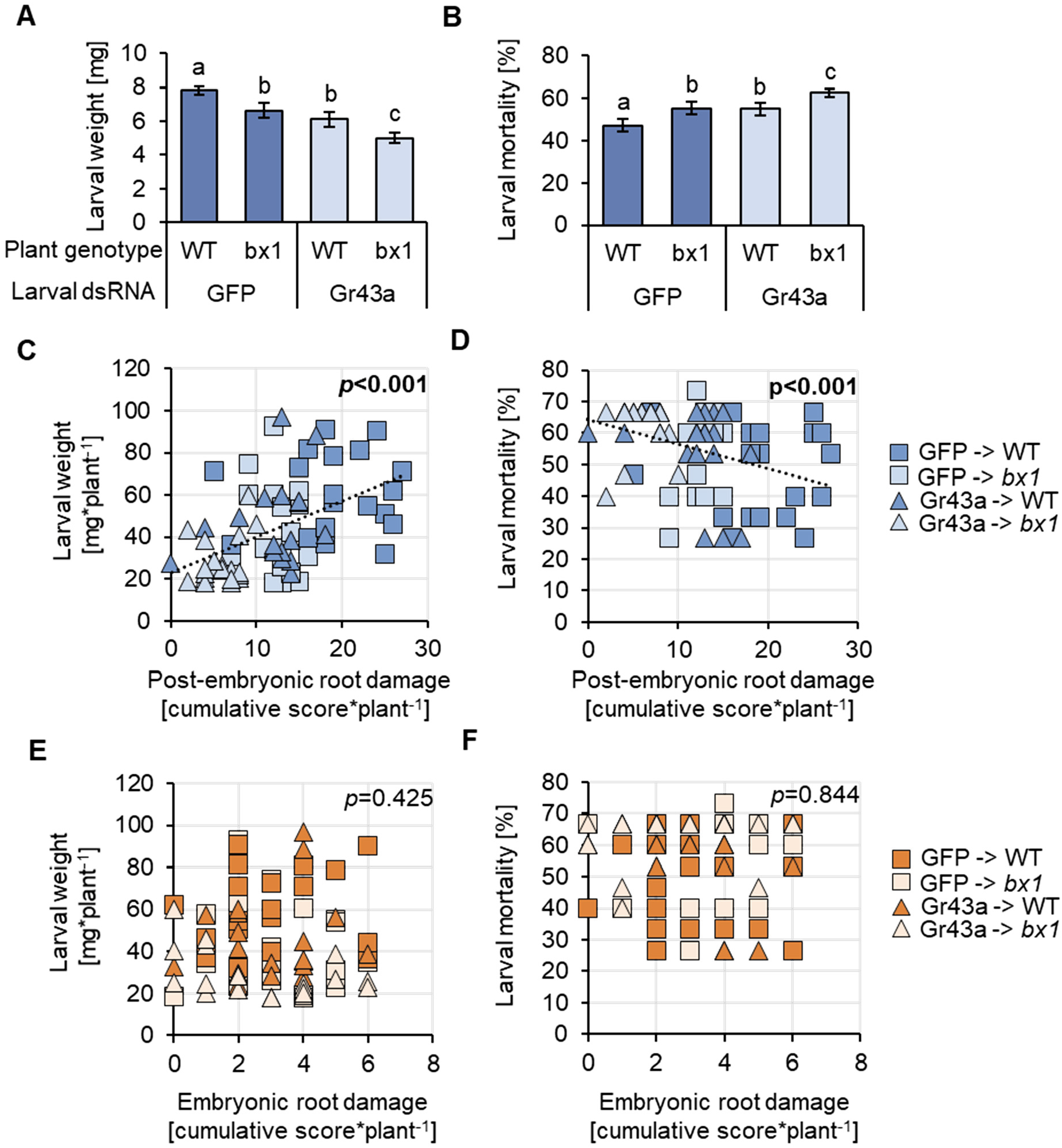
Using benzoxazinoids and sugars as foraging cues improves herbivore growth and survival. (*A*) Larval weight of control (GFP) and DvvGr43a-silenced dsRNA fed western corn rootworm larvae feeding on wild type B73 or *bx1* mutant plants for 7 days. Note that in a no-choice situation, neither the *bx1* mutation nor DvvGr43a silencing reduce larval performance ((69) and Fig. S5). Letters indicate significant differences (p<0.05, Holm-Sidak post hoc tests, n=20 pots with 15 larvae each). (*B*) Larval mortality within the same experiment. Letters indicate significant differences (p<0.05, Holm-Sidak post hoc tests, n=20 pots with 15 larvae each). (*C*-*F*) Correlations between cumulative damage per plant and larval performance parameters for post-embryonic roots (*C-D*) and embryonic roots (*E-F*). Linear regressions are shown for significant correlations (p<0.05). *P*-values are shown for Spearman Rank Order correlations.

## Discussion

Plant primary and secondary metabolites can act together to determine the behavior of generalist herbivores on artificial diet (33, 34). The present work expands these findings by demonstrating that primary and secondary metabolites can have distinct, overlapping, and additive roles as determinants of the behavior, and, consequently, the performance of a specialist herbivore *in planta*.

Herbivores are thought to use primary and secondary metabolites to make foraging decisions within complex host metabolomes (16), but how they combine these two types of cues during different stages of host finding and acceptance is not well understood. We find that the western corn rootworm employs sugars, but not benzoxazinoids for host finding. Sugars increase the successful localization of post-embryonic roots, but only in the presence of benzoxazinoids, which are essential for the latter. In contrast to these distinct roles, sugars and benzoxazinoids have similar roles in enhancing overall root damage and tissue-specific post-embryonic root damage, and act in an additive manner in this context. Thus, western corn rootworm larvae employ primary and secondary metabolites in varying combinations and hierarchies for different behavioral patterns. This finding differs from current hypotheses and observations for polyphagous insects that suggest sequential use of different foraging cues (84, 85). The explanation for this discrepancy is likely found in the distribution patterns of plant primary and secondary metabolites in maize roots and rhizosphere. Benzoxazinoids and sugars constantly accumulate in maize roots, with stable concentration differences between post-embryonic and embryonic roots (66, 69). Both compound classes are also exuded into the rhizosphere (86, 87). Current work using enzyme-inhibiting and non-inhibiting extraction solvents (66, 69, 86) suggests that benzoxazinoids are likely exuded as glucosides before being deglycosylated at the root surface, resulting in the release of aglucones and glucose. Aglucones such as DIMBOA then form attractive complexes with free iron. (66). Thus, both sugars and benzoxazinoids are present inside and outside of the roots, with outside accumulation likely being more dynamic than inside accumulation. Taking into account that benzoxazinoids are plant-specific, while sugars are more general cues (88, 89), one can envisage a scenario where the western corn rootworm relies on sugars as general short-distance cues for the presence of roots (with potentially high concentrations on the surface of maize roots due to benzoxazinoid deglycosylation), followed by benzoxazinoids as specific within-plant tissue selection cues. Benzoxazinoids may also be used as specific short-distance host recognition cues in case roots of multiple plant species are intermingled. Following host and tissue finding, the herbivore would then use both sugars and benzoxazinoids as stable host acceptance and feeding cues. High resolution metabolite imaging (90) together with the manipulation of gustatory receptors (23) could help to test these hypotheses and explore the connections between the distribution dynamics of different plant metabolites and herbivore foraging.

Plants produce complex chemical mixtures, and herbivores likely combine many of these chemicals to make appropriate foraging decisions (11, 12, 29). In line with this idea, recent *in vivo* work shows that, while individual compound classes and insect gustatory receptors can be of major importance for specific behavioral patterns, their disruption is not sufficient to fully suppress these behaviors (22–24). In the case of the western corn rootworm, we find that even the suppression of two major classes of foraging cues does not fully abolish its ability to find and accept a host plant, at least under laboratory conditions. Several other plant-derived chemicals, including different volatiles and fatty acids, are known to attract, arrest and stimulate feeding by the western corn rootworm (61, 62, 64). These cues are likely to enable basic behavioral patterns in the absence of sugars and benzoxazinoids. Whether the residual levels of benzoxazinoids that are still present in *bx1* mutant roots contribute to foraging success remains to be determined. *bx1.igl* double mutants that no longer produce benzoxazinoids are only available in a segregating genetic backgrounds, thus making direct comparisons difficult (81). However, under no choice-conditions, western corn rootworm larvae have been observed to attempt to feed on rice, a plant species that does not produce benzoxazinoids (66), thus demonstrating that some degree of host acceptance also occurs in the absence of these chemicals. We conclude that the flexible integration of complex blends of chemical cues provides partial resistance to chemically mediated behavioral disruption. Recent work highlights the importance of learning in herbivore foraging (12, 91), which likely increases the capacity of herbivores to respond to dynamic chemical landscapes and to substitute between different cues with redundant information content. How previous experiences influence the foraging behavior of the western corn rootworm remains to be determined. All larvae used in our experiments were reared on maize, reflecting their high degree of specialization on this host plant (59). Western corn rootworm adults are more flexible with their diet (59, 92), and it would be interesting to understand how their experience influences the responses of the next larval generation to host plant cues.

The capacity of an herbivore to select a host plant and feed on specific tissues generally improves its fitness (40, 42, 93). In line with these preference-performance relationships, diet experiments often document strong associations between chemically-mediated herbivore feeding preferences and herbivore performance (17). In many of these cases, herbivore feeding is directly elicited by substances that improve performance (17). However, plant metabolomes are likely to complex to be fully assessed by an herbivore, who thus need to rely on digestive feedbacks or subsets of chemical cues to make decisions. To what extent the use of plant chemical cues enhances herbivore performance by guiding herbivores to nutritious tissues is poorly understood, and interactions between primary and secondary metabolite chemical cues have, to the best of our knowledge, not been tested *in planta*. Our experimental approach rules out direct effects of benzoxazinoids and sugar perception on western corn rootworm performance (66, 69) and thus allows us to quantify their net impact as foraging cues. We find that the availability of sugars and benzoxazinoids during foraging improves herbivore weight gain and survival individually by 8%, and in combination by 15%, benefits which are most likely driven by increased post-embryonic root feeding. Post-embryonic roots of young maize plants are growing more vigorously than embryonic roots, receive a higher share of photosynthates and are richer in carbohydrates, amino acids and total soluble protein and increase western corn rootworm growth in no-choice experiments (69). In combination, these findings demonstrate that the combined use of primary and secondary metabolites as foraging cues enhance the performance of a specialist insect herbivore by guiding it to its most profitable feeding niche.

The gustatory receptor Gr43a is a highly conserved insect sugar receptor that is narrowly tuned to D-fructose (49), but mediates feeding preferences for different sugars through dietary feedbacks (55). Given the conserved function, predicted protein structure, tissue-specific expression pattern and effect on sugar preferences, *DvvGr43a* is very likely to act as a sugar sensor in the western corn rootworm as well. An interesting observation in this context is that *DvvGr43a*-dependent larval feeding preferences for sugars and sugar mixtures are only established after 2-3 hours, while feeding preferences for Fe(III)(DIMBOA)3 are established more rapidly (66). In Drosophila, Gr43a increases sugar intake under food limiting conditions and suppresses sugar intake under satiation (49). We thus hypothesized that the delayed *DvvGr43a*-dependent sugar feeding preference of western corn rootworm larvae for sugars may be the result of a starvation response. However, we observed a similar lag time in sugar preference in pre-starved larvae. This suggests that *DvvGr43a* likely functions as a sugar sensor rather than as a nutrient uptake regulator in the western corn rootworm. The observed lag phase may be explained by the potential role of *DvvGr43a* as an internal sugar sensor that triggers behavioral responses once sugars are taken up and elicit changes in hemolymph sugar levels. A deeper molecular characterization could shed light on the precise neurological and physiological roles of *DvvGr43a* in the western corn rootworm in the future.

In conclusion, the experiments presented here reveal how primary and secondary metabolites interact to guide the foraging of a specialist herbivore. The distinct individual and combined roles of these metabolites reveal finely tuned herbivore foraging strategies that are directly linked to improved herbivore performance and survival. At the same time, foraging is remarkably robust to perturbation, most likely due to the flexible use of redundant host plant cues. Our mechanistic approach thus offers insights into the complex, multilayered interplay between plant metabolomes and chemically guided herbivore foraging.

## Materials and Methods

### Plants and insects

Maize seeds (*Zea mays* L., inbred line B73) were provided by Delley Semences et Plantes SA (Delley, CH). The near-isogenic benzoxazinoid deficient *bx1* mutant line in a B73 background was obtained by backcrossing the original *bx1* mutant five times into B73 (94). Seedlings were grown under greenhouse conditions (23 ± 2°C, 60% relative humidity, 16:8 h L/D, and 250 mmol/m^2^/s^1^ additional light supplied by sodium lamps). Plantaaktiv^®^ 16+6+26 Typ K fertilizer (Hauert HBG Dünger AG, Grossaffoltern, Switzerland) was added twice a week after plant emergence following the manufacturer’s recommendations. When plants were used to feed insects, seedlings were germinated in vermiculite (particle size: 2-4 mm; tabaksamen, Switzerland) and used within four days after germination. Eggs of the non-diapausing *Diabrotica virgifera virgifera* strain were originally supplied by USDA-ARS-NCARL, Brookings, SD. Insect colonies were subsequently established and are maintained in the University of Bern and the University of Neuchatel. Upon egg hatching, insects were maintained in organic soil (Selmaterra, Bigler Samen AG, Thun, Switzerland) and fed freshly germinated maize seedlings.

### Herbivore behavior

To evaluate the preference of western corn rootworm larvae for different plant genotypes and root types, we use three different experimental setups: potted plants, root systems laid out in petri dishes and detached roots in petri dishes.

For western corn rootworm feeding preference experiments using whole plants, three-week-old maize plants grown in 200 mL cylindrical plastic pots filled with sand and topped off with potting soil were infested with fifteen second instar western corn rootworm larvae. Twenty plants per plant genotype/type of larvae combination were evaluated (n=20). Seven days after infestation, plants were gently excavated and inspected for root damage. Damage was recorded for each root individually using the following scale. 0: no visual damage, 1: one-three bite marks, 2: more than three bite marks, 3: several bite marks and brown discoloration, 4: root partially removed, and 5: root fully removed.

Western corn rootworm feeding preference assays using petri dishes were conducted as described (95, 96). Briefly, two-week-old maize plants grown as described above were gently excavated. The root systems were washed and laid out onto a moist filter paper embedded in a petri dish (14 cm diameter, Greiner Bio-one, Austria). A circular whole in the rim of the dishes accommodated the plant stems, with the shoots remaining outside of the dishes. Five third instar western corn rootworm larvae were released into each petri dish. Petri dishes were sealed and then covered with aluminum foil. The number of western corn rootworm larvae feeding on each root type was determined at different time points after their release.

To evaluate western corn rootworm feeding preference using detached roots, pieces of equal sizes of two post-embryonic roots and two embryonic roots from two-week-old maize plants were placed next to each other onto a moist filter paper (diameter 90 mm, GE Healthcare, UK) embedded into a petri plate (90 mm diameter, Greiner Bio-one, Austria). Five third instar western corn rootworm larvae were released into each petri dish. Petri plates were sealed and then covered with aluminum foil. The number of western corn rootworm larvae feeding on each root type was determined at different time points.

To evaluate the attractiveness of CO_2_ for western corn rootworm larvae, we conducted choice experiments using belowground olfactometers (64). CO_2_ levels were increased in one arm of the olfactometer by delivering CO_2_-enriched synthetic air (1% CO_2_, Carbagas, Switzerland). Olfactometer arms were closed on top using parafilm during CO_2_ delivery. A manometer connected to the synthetic air bottle allowed to fine-tune CO_2_ delivery rates. CO_2_ levels were measured using a gas analyzer (Li7000, Li-Cor Inc., Lincoln, Nebraska, USA). Once desired CO_2_ concentrations were reached, CO_2_ delivery was terminated, olfactometer arms were connected to olfactometer central connectors and six second to third instar western corn rootworm larvae were release immediately and their choice were evaluated within ten minutes of release. Six olfactometers per larval type were assayed (n=6).

To evaluate the attractive effects of Fe(III)(DIMBOA)_3_ for western corn rootworm larvae, we release six second to third instar larvae in the middle of a moist sand-filled petri dish (9 cm diameter, Greiner Bio-One GmbH, Frickenhausen, DE) where they encountered a filter paper disc treated with 10 μl of Fe(III)(DIMBOA)3 (1 μg/ml of water) or, on the opposite side, a filter paper disc treated with water only. Ten independent petri dishes per larval type were evaluated (n=10). Fe(III)(DIMBOA)3 was prepared fresh by mixing FeCl3 and DIMBOA at a 1:2 ratio as described (96). Larval preferences were recorded 1 h after releasing the larvae.

To evaluate attractive effects of soluble sugars for western corn rootworm larvae, we followed the procedure described by 97 with minor modifications. Briefly, we released six second to third instar western corn rootworm larvae in the middle of a moist sand-filled petri dish (6 cm diameter, Greiner Bio-One GmbH, Frickenhausen, DE) where they encountered a filter paper disc treated with 10 μl of either glucose (30 mg/ml), fructose (30 mg/ml), sucrose (30 mg/ml), or a mixture of the three (10 μl containing each sugar at a concentration of 30 mg/ml), or, on the opposite side, a filter paper treated with water only. Fifteen independent petri dishes per treatment were assayed (n=15). Larval preferences were recorded regularly for 3 h. In a second experiment, larvae were pre-fed on maize roots or starved for 12 h before being released into petri dishes with filter paper discs treated with a sugar mixture or water, as described above. Eight independent petri dishes with six larvae each were assayed (n=8).

### Herbivore performance

To evaluate the impact of benzoxazinoids and sugars on western corn rootworm performance, three-week-old maize plants grown in 200 mL cylindric plastic pots (11 cm depth and 4 cm diameter) filled with sand were infested with fifteen second instar western corn rootworm larvae (n=20). Seven days after infestation, the larvae were collected, counted, and weighted using a microbalance. Note that weight and survival was determined on the same plants as root damage (see section “Herbivore behavior”).

To evaluate the impact of *DvvGr43a*-silencing on western corn rootworm performance in a no-choice setting, GFP and *DvvGr43a* fed first instar larvae were reared on 3-4-day old maize seedlings, which only produce embryonic roots, in soil-filled plastic cups (n=40). Nine larvae were used per cup. Fresh seedlings were provided to the larvae every three days for a total of six days, after which the larvae were collected, counted, and weighted using a microbalance.

### Root metabolite measurements

Untargeted root metabolomics was conducted as described (81). Embryonic and post-embryonic roots of B73 plants were harvested separately, flash-frozen and ground in liquid nitrogen. Sixty to eighty mg of plant tissues were then extracted in 800 μL of acidified H2O/MeOH (50:50 v/v; 0.1% formic acid). Resulting root extracts were analysed using an Acquity UHPLC system coupled to a G2-XS QTOF mass spectrometer equipped with an electrospray source (Waters). Gradient elution was performed on an Acquity BEH C18 column (2.1 × 50 mm i.d., 1.7 μm particle size) at 1-27.5% B over 3.5 min, 27.5-100% B over 1 min, holding at 100% B for 1 min, and reconditioning at 1% B for 1 min, where A =0.1% formic acid/water and B = 0.1% formic acid/acetonitrile. The flow rate was 0.4 mL/min. The temperature of the column was maintained at 40°C, and the injection volume was 1 μL. The QTOF MS was operated in negative mode. The data were acquired over an m/z range of 100–1200 with scans of 0.15 s at collision energy of 4 V and 0.2 s with a collision energy ramp from 10 to 40 V. The capillary and cone voltages were set to 2 kV and 20 V, respectively. The source temperature was maintained at 140°C, the desolvation gas was 400°C at 1000 L h-1 and cone gas flow was 50 L/hr. Accurate mass measurements (<2 ppm) were obtained by infusing a solution of leucin encephalin at 200 ng/mL at a flow rate of 10 μL/min through the Lock Spray probe (Waters). The chromatograms were processed using Progenesis QI (Nonlinear Dynamics, Newcastle UK) with default settings for spectral alignment, peak picking, and deconvolution, resulting in a total of 4956 mass features. Of these, 444 mass features eluting between 0 and 0.3 min were discarded, since their lack of retention and excessive coelution made reliable identification impossible.

Abundances for each remaining mass feature were normalised by ArcSin transformation, and mean normalized abundances for post-embryonic and embryonic roots were compared by Student’s t-test. P-values were adjusted by Benjamini–Hochberg false discovery rate multiple testing correction and a fold-change cut-off of 2.0 was applied to generate a list of compounds that significantly differed between treatments. Volcano plots were used to visualize treatment differences, and the list of differentially expressed features was tested against an in-house mass-fragmentation database using Progenesis QI (Progenesis MetaScope v1.0.6253.26544, Nonlinear Dynamics, Newcastle, UK). In addition, chromatograms were manually scanned for exact masses of known plant metabolites, specifically amino acids, sugars, organic acids, and flavonols. Potential matches were likewise tested against the mass fragmentation database, and putative identifications were accepted based on exact mass, mass fragmentation patterns, and relative retention times.

Root benzoxazinoid quantification was conducted as described (98). For this, embryonic and post-embryonic roots of B73 or *bx1* plants were harvested separately, flash-frozen and ground in liquid nitrogen. Approximately 100mg of plant tissues were then extracted in 1 mL of acidified H2O/MeOH (50:50 v/v; 0.1% formic acid), and analyzed with an Acquity UHPLC system coupled to a UV detector and a Waters QDa mass spectrometer equipped with an electrospray source(Waters). Compounds were separated on an Acquity BEH C18 column (2.1×50 mm i.d., 1.7 μm particle size). Water (0.1% formic acid) and acetonitrile (0.1% formic acid) were employed as mobile phases A and B. The elution profile was: 0-1 min, 10% B; 1-4 min, 10-30% B; 4-5 min, 30-9740% B; 5-6 min, 40-100% B; 6-8.5 min, 100% B; followed by 1.5 min reconditioning at 10% B. The mobile phase flow rate was 0.4 mL/min. The column temperature was maintained at 40°C, and the injection volume was 3 μL. The MS was operated in negative mode and data were acquired in single ion recording (SIR) mode, using [M-H]^-^ and [M+HCOO^-^]^-^ adducts of benzoxazinoid compounds. All other MS parameters were left at their default values as suggested by the manufacturer. Benzoxazinoid peaks were identified from MS signals and quantified from integrated UV signals at 265nm. External standards for HMBOA, DIMBOA, DIMBOA-Glc, HDMBOA-Glc, and MBOA were used for absolute quantification. Where concentrations were too low to result in reliable UV signals, a compound-specific MS-to-UV signal conversion factor was estimated to calculate ‘predicted’ UV signals from the more sensitive MS signals before quantification.

Root sugar levels were measured by an enzymatic/spectrophotometric method as described (99). Embryonic and post-embryonic roots of B73 and *bx1* plants were harvested separately, flash-frozen and ground in liquid nitrogen. Soluble sugars were extracted from 100 mg root tissue using 80% (v/v) ethanol, followed by an incubation step (10 min at 78°C) with constant shaking at 800 rpm. Pellets were re-extracted twice with 50% (v/v) ethanol (10 min at 78°C with constant shaking at 800 rpm). Supernatants from all extraction steps were pooled together, and sucrose, glucose and fructose were quantified as described (100).

### Identification of *DvvGr43a*

To identify the *Gr43a* receptor orthologues in the western corn rootworm, we used the *Gr43a* receptor-encoding gene sequence of *Drosophila melanogaster* as query against publicly available nucleotide sequences and against the western corn rootworm genome sequences (NCBI accession: PXJM00000000.2) using the National Center for Biotechnology Information Basic Local Alignment Search Tool (NCBI BLAST). The full gene sequence can be retrieved from the National Center for Biotechnology Information (NCBI) data bank using the following accession number: XM_028279548.1. Then, we reconstructed evolutionary relationships between the identified DvvGr43a gene and several other putative Gr43a genes that have been functionally characterized as glucose, fructose, and/or sucrose receptors (49, 50, 53, 54, 75–78). The evolutionary relationships were inferred using the Maximum Likelihood method based on the Le Gascuel 2008 model in MEGA7 (101, 102). The tree with the highest log likelihood (−14287.71) is shown. The percentage of trees in which the associated taxa clustered together is shown next to the branches. Initial tree(s) for the heuristic search were obtained automatically by applying Neighbor-Join and BioNJ algorithms to a matrix of pairwise distances estimated using a JTT model, and then selecting the topology with superior log likelihood value. A discrete Gamma distribution was used to model evolutionary rate differences among sites (5 categories (+G, parameter = 1.0811)). The tree is drawn to scale, with branch lengths measured in the number of substitutions per site. There was a total of 569 positions in the final dataset. Graphical representation and edition of the phylogenetic tree were performed with the Interactive Tree of Life (version 3.5.1) (103). Protein tertiary structures and topologies were predicted using Phyre2 (104). The protein sequences used to reconstruct evolutionary relationships can be retrieved from the NCBI under the following accession numbers: *Heliothis virescens* Gr4 (CAD31946.1), *Danaus plexippus* Gr4 (EHJ77681.1), *Bombyx mori* Gr9(NP_001124345.1), *Bombus impatiens* Gr43a (XP_003486787.1), *Nasonia vitripennis* Gr3 (NP_001164386.1), *Apis mellifera* Gr3 (XP_016768876.1), *Drosophila melanogaster* Gr43a (NP_523650.2), *Tribolium castaneum* Gr20 (EFA05758.1), *Anopheles gambiae* Gr43a (XP_318100.4), *Camponotus floridanus* Gr43a (EFN61344.1), *Aedes aegypti* Gr43a (XP_001658898.3), *Culex quinquefasciatus* Gr43a (XP_001842305.1), *Orussus abietinus* Gr43a (XP_012270798.1), *Aethina tumida* Gr43a (XP_019876958.1), *Anoplophora glabripennis* Gr43a (XP_018574194.1), *Helicoverpa armigera* Gr4 (AGK90011.1), *Helicoverpa armigera* Gr9 (JX970522.1), *Trichogramma chilonis* Gr43a (QAY30709.1), and *Diabrotica virgifera* Gr43a (XP_028135349.1). For *Plutella xylostella* GR43a-1 and *Plutella xylostella* GR43a-2, and *D. melanogaster* gustatory receptors refer to: (53, 105), as these sequences have not been deposited in a public repository.

### Double stranded RNA production

Double stranded RNA targeting *DvvGr43a* was produced by *Escherichia coli* HT115. To this end, electrocompetent *E. coli* HT115 cells were transformed with recombinant L4440 plasmids that contained either a 240 bp long *DvvGr43a* gene fragment, or a 240 bp long green fluorescent protein (GFP) gene fragment flanked by two T7 promotors in opposite directions. Gene fragments were synthetized *de novo* (Eurofins, Germany). To induce the production of dsRNA, an overnight bacterial culture was used to inoculate fresh Luria-Berthani broth (25 g/L, Luria/Miller, Carl Roth GmbH, Karlsruhe, Germany). Once the bacterial culture reached an OD600 of 0.6-0.8, it was supplemented with isopropyl β-D-1-thiogalactopyranoside, IPTG) (Sigma-aldrich, Switzerland) at a final concentration of 2mM. Bacterial cultures were incubated at 37°C in an orbital shaker at 130 rpm for 16h. Bacteria were harvested by centrifugation and stored at −20°C for further use (106).

### Gene silencing experiments

To induce gene silencing in western corn rootworm, six to ten western corn rootworm larvae were released in solo cups (30 mL, Frontier Scientific Services, Inc., Germany) containing approx. 2g of autoclaved soil (Selmaterra, Bigler Samen AG, Thun, Switzerland) and 2-3 freshly germinated maize seedling. Maize seedlings were coated with 1 ml of bacterial solution containing approximately 200-500 ng of dsRNA targeting *DvvGr43a*. As controls, larvae were fed with bacteria producing dsRNA targeting green fluorescent protein genes (GFP). Seedlings coated with bacteria were provided every other day for three consecutive times. Two days after, larvae were collected and used for experiments.

### Gene expression measurements

Total RNA was isolated from approximately 10 mg of frozen, ground and homogenized western corn rootworm larval tissue (3-7 larvae per biological replicate, n=8-11) using the GenElute Universal Total RNA Purification Kit (Sigma-Aldrich, St. Louis, MO, USA). A NanoDrop spectrophotometer (ND-1000, Thermo Fisher Scientific, Waltham, MA, USA) was used to estimate RNA purity and quantity. DNase-treated RNA was used as template for reverse transcription and first strand cDNA synthesis with PrimeScript Reverse Transcriptase (Takara Bio Inc., Kusatsu, Japan). DNase treatment was carried out using the gDNA Eraser (Perfect Real Time) following manufacturer’s instructions (Takara Bio Inc.). For gene expression analysis, 2 μl of undiluted cDNA (i.e. the equivalent of 100 ng total RNA) served as template in a 20 μl qRT-PCR using the TB Green Premix Ex Taq II (Tli RNaseH Plus) kit (Takara Bio Inc.) and the Roche LightCycler 96 system (Roche, Basel, Switzerland), according to manufacturer’s instructions. Transcript abundance of the following western corn rootworm genes were analysed: *DvvGr43a* and *DvvGr2. Actin* was used as reference gene to normalize expression data across samples. Relative gene expression levels were calculated by the 2^-ΔΔCt^ method (107). The following primers were used: DvvGr43a-F GTCACATTCACCACGGGTCT, DvvGr43a-R CGTTCGGTCTCTAACTTTGGC, DvvGr2-F GAACTAAGCGAGCTCCTCCA, DvvGr2-R CAGAAGCACCATGCAATACG, DvvActin-F TCCAGGCTGTACTCTCCTTG, and DvvActin-R CAAGTCCAAACGAAGGATTG.

### Statistical analyses

Differences in root benzoxazinoid and root sugar levels as well as gene expression levels were analyzed by One- and Two-Way Analyses of Variance (ANOVAs). Normality and equality of variance were verified using Shapiro–Wilk and Levene’s tests, respectively. Holm–Sidak post hoc tests were used for multiple comparisons. Datasets from experiments that did not fulfill the assumptions for ANOVA were natural log-, root square-, or rank-transformed before analysis. Differences in damage scores between embryonic and post-embryonic roots were assessed using Wilcoxon Signed Rank Tests. Differences in average root damage scores between treatments were assessed using Kruskal-Wallis ANOVAs on ranks followed by Tukey’s post hoc tests. The above analyses were performed in Sigma Plot 12.0 (SystatSoftware Inc., San Jose, CA, USA) with the support of the built-in Advisor Wizard function. Differences in larval preference were assessed through Generalized Linear Models (GLM) with binomial distribution and corrected for overdispersion with quasi-binomial function when necessary followed by analysis of variance (ANOVA) and FDR-corrected Least Square Means post hoc tests. These analyses were followed by residual analysis to verify the suitability of the error distribution and model fitting. These Analyses were conducted using R 3.2.2 (43) using the packages “lme4”, “car”, “lsmeans” and “RVAideMemoire” (108–112). Mass features were normalized using the function *arcsinh_x* in the R package “bestNormalize” and compared between root types using functions *t.test* in base R and *foldchange* in the package “gtools”. P-values were FDR-corrected using the function *p.adjust*.

## Acknowledgements

We thank Elisa J. Bernklau (Colorado State University) for advice on sugar preference experiments.

## Funding

This project was supported by a European Union Horizon 2020 Marie Sklodowska-Curie Action (MSCA) Individual Fellowship (Grant # 794947 to B.C.J.S.) and the Swiss National Science Foundation (Grants # 155781, 160786 and 157884 to M.E.).

## Competing interests

The Authors declare no competing interests.

## Data and materials availability

The data generated for this manuscript is available on Dryad [DOI to be inserted].

---

**Table S1.** Identified metabolites in maize post-embryonic and embryonic roots. Metabolites with retention times highlighted by asterisks (*) eluted together with many weakly-retained compounds that could not be separated by reversed-phase liquid chromatography (RP-UHPLC). Their identification is therefore tentative and based on accurate masses only.

| Nr | Compound class | Compound Name | Molecular formula | Retent ion time [min] | MS adducts | Progenes is QI fragment ation score | Characteristic fragments [m/z] |
| --- | --- | --- | --- | --- | --- | --- | --- |
| 1 | Benzoxazinoid | DHBOA-Glc | C<sub>14</sub>H<sub>17</sub>NO<sub>9</sub> | 1.30 | M-H, M+FA-H, 2M-H | 16.5 | 180.0303, 162.0200 |
| 2 | Benzoxazinoid | DIBOA-Glc | C<sub>14</sub>H<sub>17</sub>NO<sub>9</sub> | 1.91 | M-H, M+FA-H, 2M-H | 2.6 | 180.0303, 162.0200, 134.0247 |
| 3 | Benzoxazinoid | HMBOA-Glc | C<sub>15</sub>H<sub>19</sub>NO<sub>9</sub> | 2.11 | M-H, M+FA-H, 2M-H | 10.3 | 194.0459, 166.0511 |
| 4 | Benzoxazinoid | DIMBOA-Glc | C<sub>15</sub>H<sub>19</sub>NO<sub>10</sub> | 2.18 | M-H, M+FA-H, 2M-H | 5.9 | 210.0406, 192.0302, 164.0356, 149.0119 |
| 5 | Benzoxazinoid | DIM<sub>2</sub>BOA-Glc | C<sub>16</sub>H<sub>21</sub>NO<sub>11</sub> | 2.22 | M-H, M+FA-H, 2M-H | 21.7 | 222.0408, 194.0458 |
| 6 | Benzoxazinoid | HMBOA | C<sub>9</sub>H<sub>9</sub>NO<sub>4</sub> | 2.30 | M-H | n/a | - |
| 7 | Benzoxazinoid | DIMBOA | C<sub>9</sub>H<sub>9</sub>NO<sub>5</sub> | 2.42 | M-H | n/a | 164.0353, 149.0118 |
| 8 | Benzoxazinoid | HDMBOA-Glc | C<sub>16</sub>H<sub>21</sub>NO<sub>10</sub> | 2.74 | M-H, M+FA-H, 2M-H | 11.5 | 224.0564, 194.0460 |
| 9 | Benzoxazinoid | HDM<sub>2</sub>BOA-Glc | C<sub>17</sub>H<sub>23</sub>NO<sub>11</sub> | 2.74 | M+FA-H | n/a | 254.0789 |
| 10 | Organic acid | Chlorogenic acid | C<sub>16</sub>H<sub>18</sub>O<sub>9</sub> | 1.77 | M-H, 2M-H | 24.3 | 191.0<br>562 |
| 11 | Organic acid | Feruloylglucose | C<sub>16</sub>H<sub>20</sub>O<sub>9</sub> | 2.05 | M-H | 81.0 | 175.0<br>404 |
| 12 | Organic acid | Sinapoylglucose | C<sub>17</sub>H<sub>22</sub>O<sub>10</sub> | 2.09 | M-H | 36.7 | 223.0615, 205.0507 |
| 13 | Organic acid | Quinic acid | C<sub>17</sub>H<sub>12</sub>O<sub>6</sub> | 0.33* | M-H, 2M-H | 70.1 | - |
| 14 | Organic acid | Coumaroylquinic acid isomer | C<sub>16</sub>H<sub>18</sub>O<sub>8</sub> | 1.70 | M-H | 98.0 | 163.0409 |
| 15 | Organic acid | Coumaroylquinic acid isomer | C<sub>16</sub>H<sub>18</sub>O<sub>8</sub> | 2.13 | M-H | 51.6 | 191.0561, 173.0455 |
| 16 | Organic acid | Feruloylquinic acid isomer | C<sub>17</sub>H<sub>20</sub>O<sub>9</sub> | 2.35 | M-H | 46.1 | 191.0562 |
| 17 | Organic acid | Feruloylquinic acid isomer | C<sub>17</sub>H<sub>20</sub>O<sub>9</sub> | 2.61 | M-H | n/a | - |
| 18 | Flavonol | Quercetin rhamnosylglucoside | C<sub>27</sub>H<sub>30</sub>O<sub>16</sub> | 2.66 | M-H | 65.7 | 301.0353 |
| 19 | Flavonol | Quercetin glucoside | C<sub>21</sub>H<sub>20</sub>O<sub>12</sub> | 2.75 | M-H | 26.0 | 301.0347, 271.0252 |
| 20 | Flavonol | Quercetin malonylglucoside | C<sub>24</sub>H<sub>22</sub>O<sub>15</sub> | 2.91 | M-H | 75.1 | 505.0986, 301.0345 |
| 21 | Flavonol | Kaempferol<br>rhamnosylglucoside | $C_{27}H_{30}O_{15}$ | 2.94 | M-H | 98.9 | 285.0412 |
| 22 | Flavonol | Kaempferol<br>glucoside | $C_{21}H_{20}O_{11}$ | 3.04 | M-H | 88.6 | 285.0395, 255.0304 |
| 23 | Flavonol | Kaempferol<br>malonylglucoside | $C_{24}H_{22}O_{14}$ | 3.25 | M-H | 96.5 | 489.1037, 285.0400 |
| 24 | Amino acid | Leucine | $C_6H_{13}NO_2$ | 0.80 | M-H | n/a | - |
| 25 | Amino acid | Phenylalanine | $C_9H_{11}NO_2$ | 1.10 | M-H | n/a | - |
| 26 | Amino acid | Tryptophan | $C_{11}H_{12}N_2O_2$ | 1.50 | M-H, M+FA-H | n/a | - |
| 27 | Sugar | Monosaccharide | $C_6H_{12}O_6$ | 0.31* | M-H | n/a | - |
| 28 | Sugar | Disaccharide | $C_{12}H_{22}O_{11}$ | 0.34* | M-H, M+FA-H,<br>2M-H | 3.2 | - |

**Table S2.** Root metabolites with behavioral activity towards western corn rootworm larvae.

| Nr | Compound class | Compound Name | Behavioral effect | Reference. |
| --- | --- | --- | --- | --- |
| 1 | Benzoxazinoid | MBOA | Host location (long-distance) | (113) |
| 2 | Benzoxazinoid | Fe-DIMBOA | Host location & acceptance (short-distance) | (66) |
| 3 | Benzoxazinoid | Fe-DIMBOA-Glc | Host location & acceptance (short-distance) | (66) |
| 4 | Sugar | Glucose | Feeding stimulation (within sugar blend) | (61) |
| 5 | Sugar | Fructose | Feeding stimulation (within sugar blend) | (61) |
| 6 | Sugar | Sucrose | Feeding stimulation (within sugar blend) | (61) |
| 7 | Fatty acid | Linoleic acid | Arrestment, feeding stimulation (within sugar blend) | (61) |
| 8 | Fatty acid | Oleic acid | Feeding stimulation (within sugar blend) | (61) |
| 9 | Fatty acid | Monoga-lactosyldiacylglycerol | Host recognition (tight turning behavior) | (62) |
| 10 | Aminobenzole | Methyl anthranilate | Host avoidance (repellent) | (63) |
| 11 | Terpene | (*E*)-β-caryophyllene | Host location and discrimination (long distance) | (43, 68) |
| 12 | Alkene | Ethylene | Host location and discrimination (long-distance) | (64) |
| 13 | Inorganic compound | CO<sub>2</sub> | Host location (long-distance) | (60) |

**Fig. S1.**
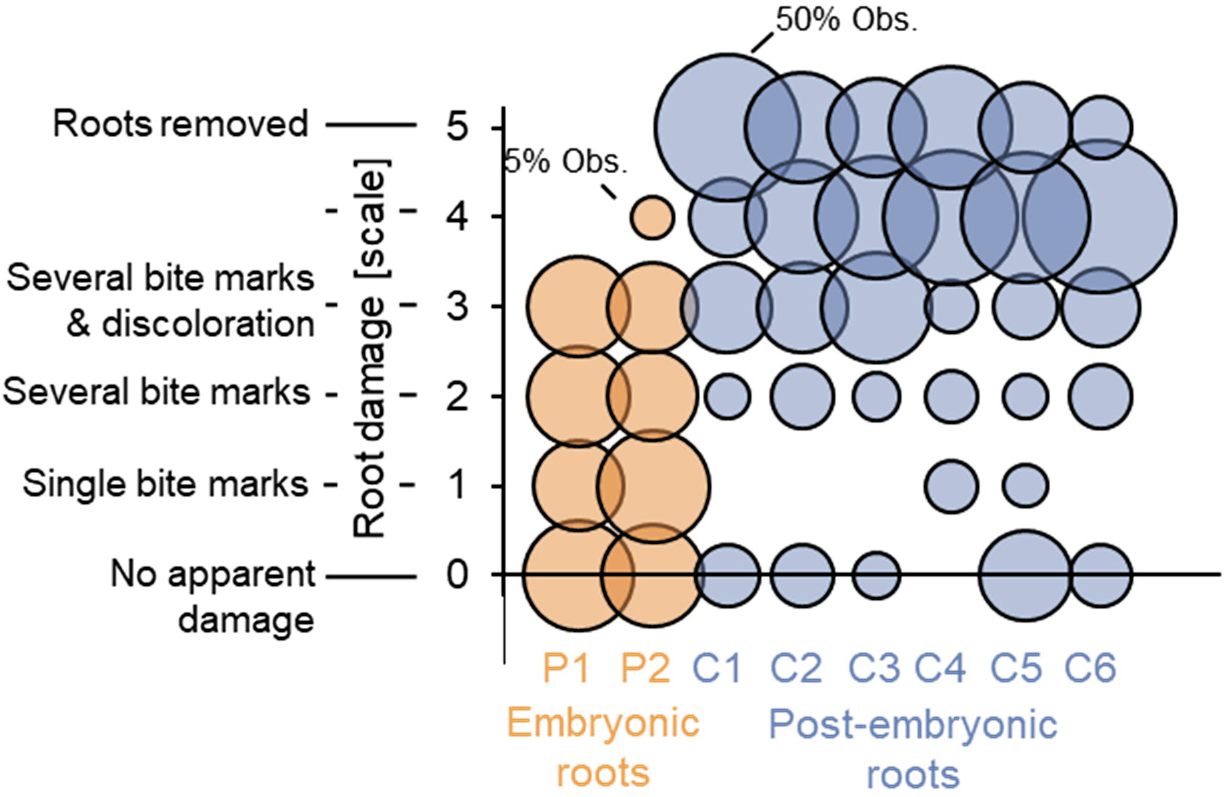
Frequency of different types of damage on individual embryonic and post embryonic roots. Data is from experiment shown in Fig. 1*B* (n=20 plants). P1-P2 refer to individual embryonic roots, C1-C6 to individual post-embryonic roots per plant. Numbers were assigned randomly to embryonic and post embryonic roots within plants. The sizes of the circles are proportional to the relative frequency (% within each root) of the different types of observed damage.

**Fig. S2.**
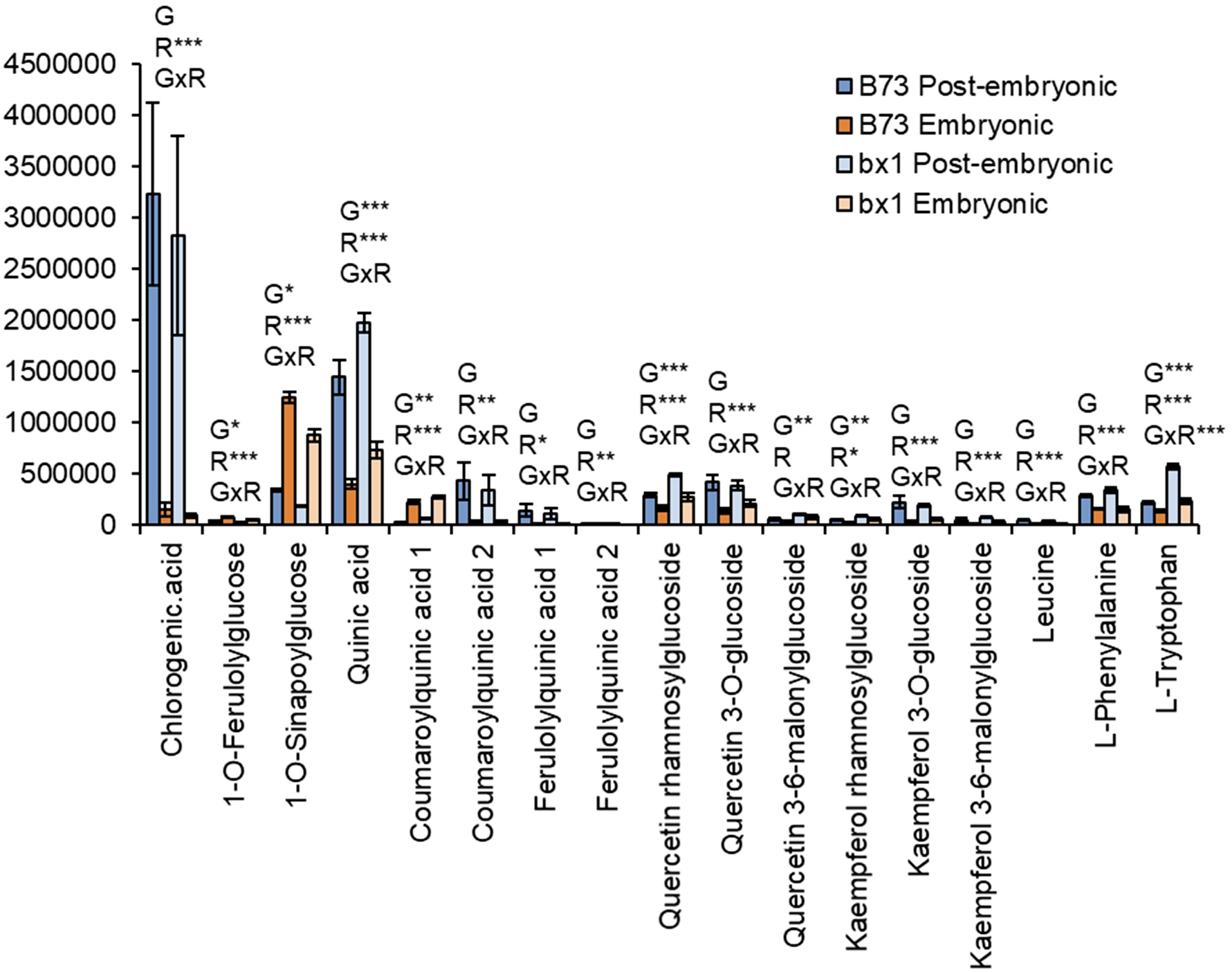
Metabolic differences between embryonic and post-embryonic roots are conserved in the bx1 mutant. Relative abundances (signal intensities) of identified metabolic features in embryonic and post-embryonic roots of wild type B73 and *bx1* mutant plants (n=7-10). For sugars and benzoxazinoids, refer to Fig. 2. Results of Two-way ANOVAs for genotype effects (G), root type effects (R) and their interaction (GxR) are shown for each compound (***p<0.001, **p<0.01, *p<0.05). Error bars denote standard errors of means (SEM).

**Fig. S3.**
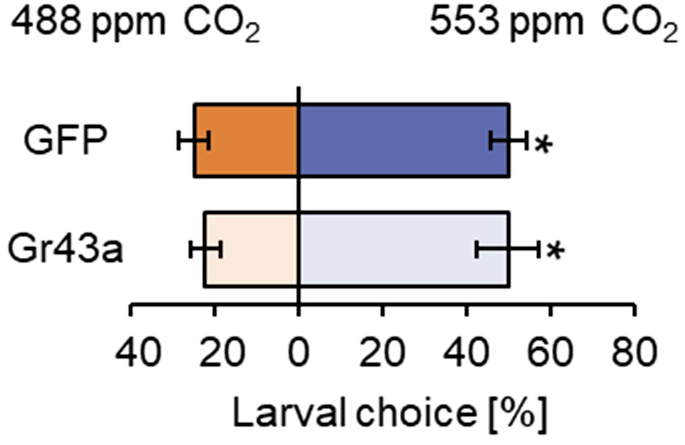
DvvGr43a does not influence the attraction of the western corn rootworm to CO_2_. Proportion of control (GFP) or *DvvGr43a*-silenced (Gr43a) larvae found on each arm of belowground olfactometers. Asterisks indicate significant differences between treatments (*p<0.05, FDR-corrected Least Square Mean post hoc tests, six two-arm olfactometers with 6 larvae each were evaluated, n=6).

**Fig. S4.**
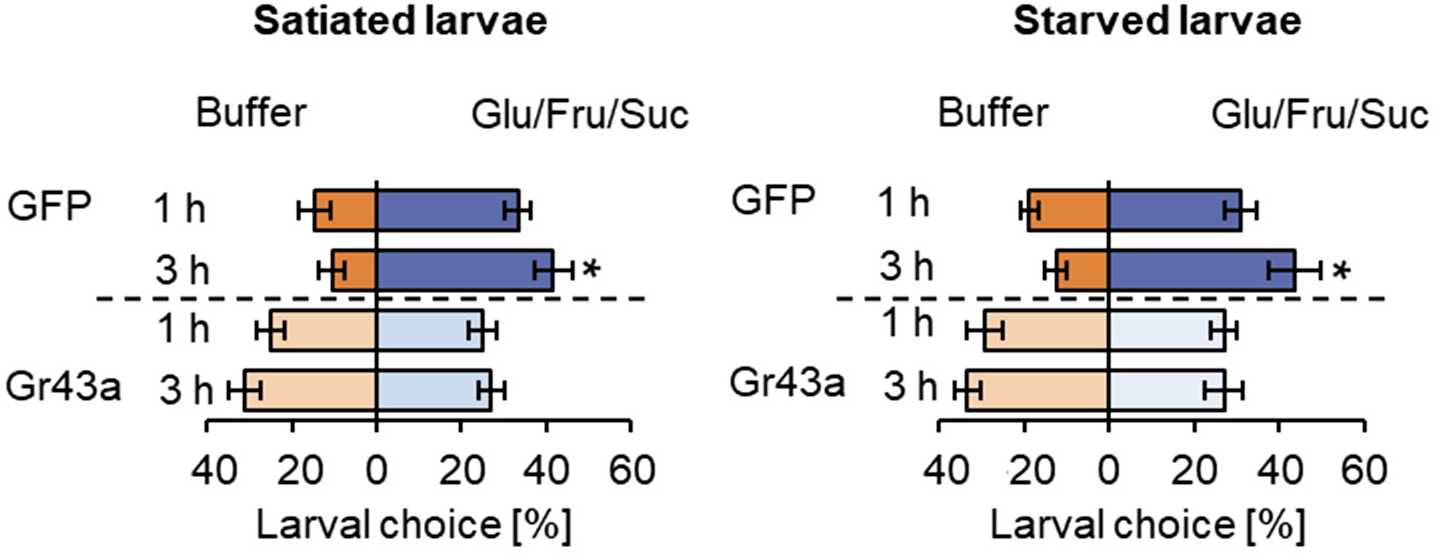
DvvGr43a mediates sugar preference independently of the feeding state of the western corn rootworm. Preference of satiated (left) and starved (right) control or *DvvGr43a*-silenced larvae for buffer or a glucose, fructose, sucrose mixture on filter discs at different time points (*p<0.05, FDR-corrected Least Square Mean post hoc tests, eight petri plates with six larvae each were assayed, n=8). Error bars denote standard errors of means (SEM).

**Fig. S5.**
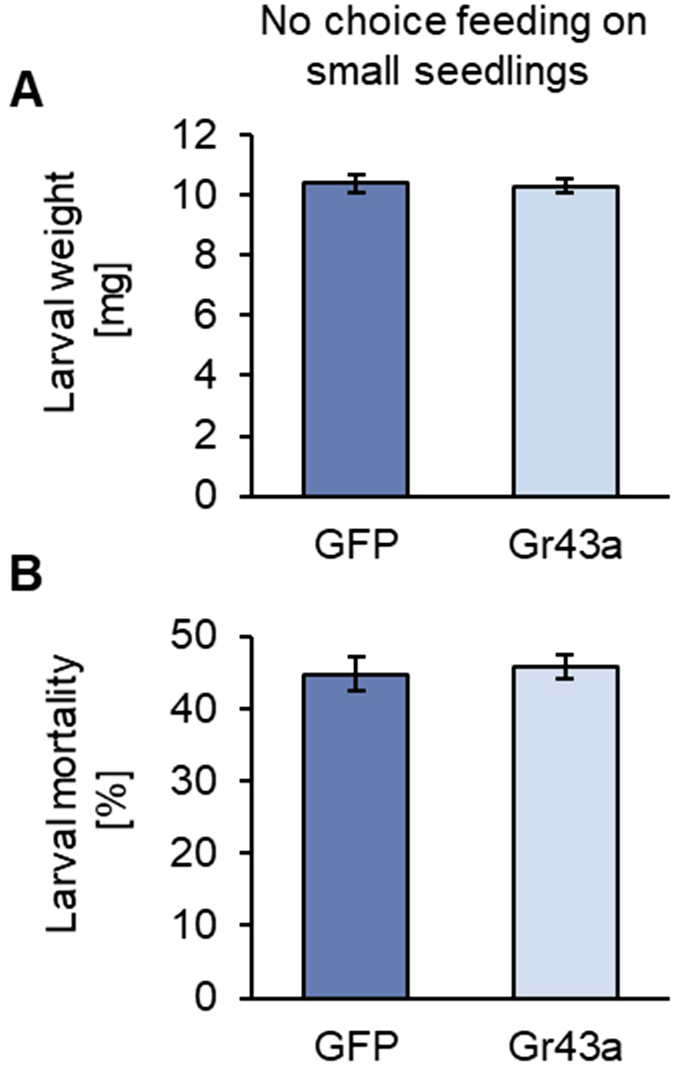
DvvGr43a does not directly impair larval performance. (*A*) Weight of western corn rootworm larvae fed on GFP and DvvGr43a dsRNA on young maize seedlings that produce embryonic roots only (no-choice setting) for 7 days (n=40 cups with 9 larvae each). (*B*) Larval mortality within the same experiment.

## Notes

### Competing Interest Statement

The authors have declared no competing interest.

### Summary of Updates

Small changes to title and abstract.

